# Micro-aggregation of a pristine grassland soil selects for bacterial and fungal communities and changes in nitrogen cycling potentials

**DOI:** 10.1101/2021.10.13.464334

**Authors:** Christoph Keuschnig, Jean M.F. Martins, Aline Navel, Pascal Simonet, Catherine Larose

**Author notes:** Address correspondence to Christoph Keuschnig.

## Abstract

Microbial analysis at the micro scale of soil is essential to the overall understanding of microbial organization and interactions, and necessary for a better understanding of soil ecosystem functioning. While bacterial communities have been extensively described, little is known about the organization of fungal communities as well as functional potentials at scales relevant to microbial interactions. Fungal and bacterial communities and changes in nitrogen cycling potentials in the pristine Rothamsted Park Grass soil (bulk soil) as well as in its particle size sub-fractions (PSFs; > 250 μm, 250-63 μm, 63-20 μm, 20-2 μm, < 2 μm and supernatant) were studied. The potential for nitrogen reduction was found elevated in bigger aggregates. The relative abundance of Basidiomycota deceased with decreasing particle size, Ascomycota showed an increase and Mucoromycota became more prominent in particles less than 20 μm.

Bacterial community structures changed below 20 μm at the scale where microbes operate. Strikingly, only members of two bacterial and one fungal phyla (Proteobacteria, Bacteroidota and Ascomycota, respectively) were washed-off the soil during fractionation and accumulated in the supernatant fraction where most of the detected bacterial genera (e.g., *Pseudomonas, Massilia, Mucilaginibacter*, *<u>Edaphobaculum</u>*, *<u>Duganella, Janthinobacterium</u>* and *Variovorax*) were previously associated with exopolysaccharide production and biofilm formation.

Overall, the applied method shows potential to study soil microbial communities at micro scales which might be useful in studies focusing on the role of specific fungal taxa in soil structure formation as well as research on how and by whom biofilm-like structures are distributed and organized in soil.

**Importance:** Intensive exploitation of soils has led to increasing environmental concerns such as pollution, erosion, emission of greenhouse gases and, in general, the weakening of its ecosystem services that are mainly regulated by microbial activity. Microbial activity and metabolism drive the formation of soil aggregates, ranging in size from a few micrometres to several millimetres. Understanding biological mechanisms related to aggregate size classes can provide insight into large-scale processes, but most research has focused on macroaggregates. Here, we investigated the microbial community and its functional changes at these smaller scales that are clearly more relevant for assessing microbial activity. We demonstrated that fungal communities are more sensitive to bigger size classes than bacteria, suggesting their dominant role in soil structure formation and turnover. We also identified preferential niches for reductive processes within the nitrogen cycle and a selection of specific taxa by analysing the water used for the wet-fractionation approach.

## Introduction

Soil harbors the highest microbial densities and diversity relative to other habitats and thus represents an important environment with a global impact on biochemical mass balances (1). The complexity and compartmentalization of soil at the microscopic scale makes it one of the most difficult microbial habitats to study (2). This compartmentalization, or soil structure, itself defined by single connected particles or so called aggregates, affects basic physical and mechanical properties such as water retention and movement, erodibility, and aeration and thus soil fertility and productivity (3). The aggregated structure of soil, in combination with the fluctuating water saturation of the available soil pore space, creates a mosaic of microenvironments varying in respiration processes (different electron acceptors). This, in turn, can generate a range of microhabitats in which specific reactions within the N-cycle might be favored. While reductive processes (denitrification) might be linked to inner parts of bigger aggregates where oxygen becomes limited, oxidative processes (ammonia oxidation) could preferentially occur on the surface regions of these aggregates (4, 5). Reactive forms of inorganic nitrogen would therefore be continuously shuttled between oxidation states in close proximity and eventually lost as N_2_O/NO gas, raising environmental concerns.

The formation, turnover and stability of aggregates depend on factors such as vegetation, soil fauna, microorganisms, cations, the interaction of clay particles and OM (6–9). A recent study highlighted the importance of microbial activity in soil structuring processes through the observation of a spatial connection between plant-derived OM and microorganisms (10). In their model of soil aggregation, Tosche et al., 2018 consider the process of microaggregate formation to be hierarchical, meaning that smaller building blocks are linked together to form larger aggregated structures that are classified by size. Here, inorganic and organic particles are considered as building units that can be assembled into composite structures by their association with OM and microorganisms such as bacteria, archaea and fungi. These composite structures can then form larger, stable microaggregates (7, 11), although no general organization of building units in aggregate structures has been found (10, 12). However, studies using soil fractionation methods to derive particle size fractions (PSFs) have shown that different physical-chemical and microbiological signatures exist among the different aggregates (13–18). These data are difficult to interpret, as the assignment of individual PSFs to specific locations in the soil matrix can be challenging, especially for microaggregates, that cannot be identified with the bare eye. These smaller particle entities could exist as composite structures, disassembled from a loosely connected bigger aggregated structure, or consist of organisms found in between the pore spaces of microaggregates.

Most soil studies address differences between macro- (> 2000 and 2000 - 250 μm) and microaggregates (< 250 μm), but few have focused on the different size classes found within microaggregates. In studies on macroaggregates, it has been shown that stabilization capacity vary among fungal taxa and are linked to specific physiological traits such as mycelial/hyphal density and enzymes production (19). In the microaggregate range less is known, although these size classes are more relevant to the individual organisms themselves given the scales at which they operate (20). Previous research has shown that most soil organisms associate with microaggregates in soil, with cells clustering either trapped in the mineral matrix, or on the outside of aggregates, as well as in the pore space (21).

An important factor in soil stabilization and aggregation is the ability to excrete extracellular polymeric substances (EPS) mainly containing polysaccharides, proteins and nucleic acids (22, 23). In addition to their physical property of acting as a glue binding soil particles, cell attachment and biofilm formation, these polymers provide crucial functions for microbes such as protection against drought, antibiotics and heavy metals, entrapment of nutrients and carbon storage, signaling, genetic exchange, adhesion and aggregation (6). Unlike bacteria, fungi are not only capable of excreting EPS and thus influencing their immediate environment, but their hyphal growth enmeshes microaggregates and stabilizes soil structures at scales up to 2 mm (24). Furthermore, a correlation between hyphal length and aggregate stability has often been observed (9, 25–27). Therefore, communities that depend on the construction of a protective biofilm structure can be expected to be located in both larger and smaller aggregates due to their hierarchical nature, whereas fungi involved in enmeshment and stabilization of macroaggregates are preferentially found in larger PSFs.

Here, we applied a gentle soil washing and separation method (Fig. 1) to disperse the soil structure and investigate microbial distribution patterns in different microaggregate structures. We hypothesized that the resulting microaggregate fractions consist of different microhabitats with distinct conditions that would spatially structure microbial communities and change nitrogen cycling potentials at the microscopic scale. We further hypothesized that fungal communities would undergo more dynamic changes over the size ranges studied, given their involvement in particles enmeshment by hyphae, while bacterial communities would not change throughout the obtained PSFs.

**Fig. 1:**
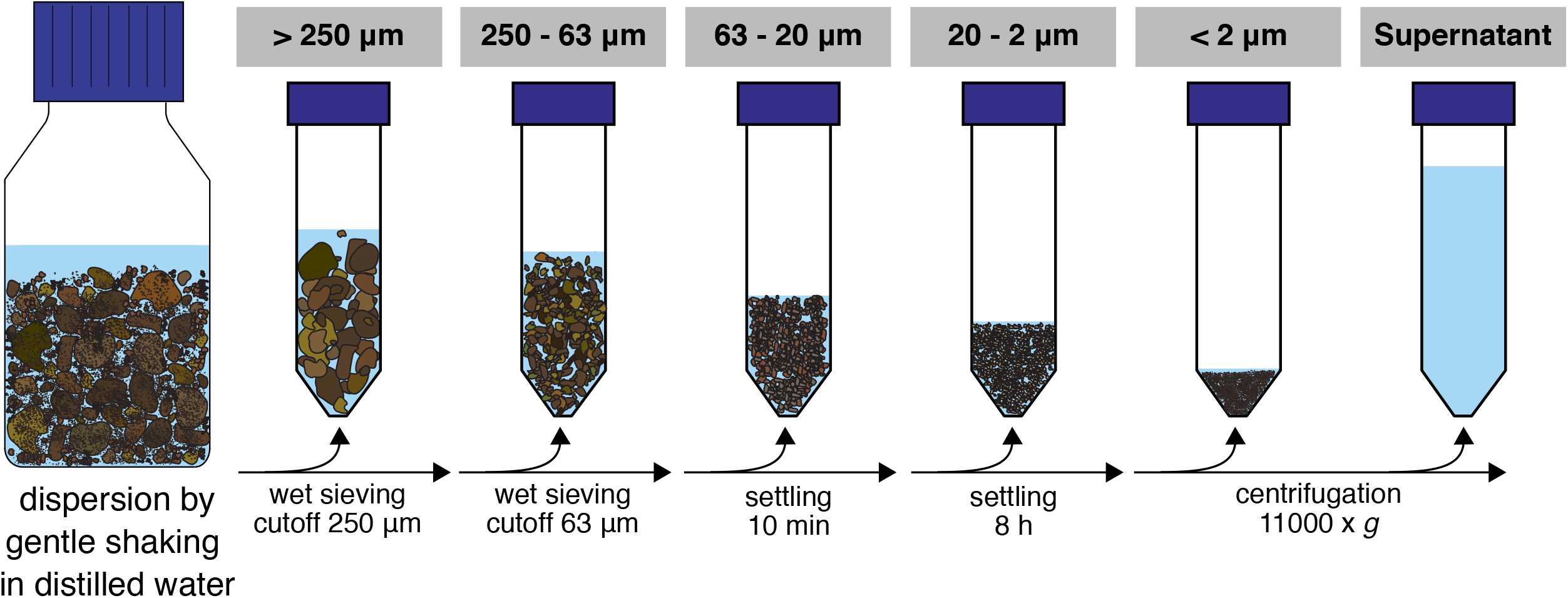
Schematic representation of the gentle soil fractionation procedure applied in the study.

## Results

### Quality of soil fractionation

The particle size distributions of the obtained fractions (Fig. 1) and bulk soil samples were analyzed by laser granulometry (Dynamic Light Scattering, DLS) to ensure that the targeted size ranges were captured (Table 1, Fig. S1). Ultrasonication on fractionated samples was used to break up stable aggregates that were not disrupted by the fractionation procedure. The distribution curves showed a shift to smaller particles with ultrasonication (US) applied in the fractions > 250 μm and 250 - 63 μm, but not below (Fig. S1). The main physical and chemical parameters obtained from the separate fractions and the bulk soil are shown in Table 1. As an estimate of the quality of the fractionation procedure, values from the single PSFs were multiplied by their mass fraction and the sum was compared to the values from bulk soil samples. While extracted DNA was overestimated with this approach, we found similar results for total carbon and nitrogen as well as NH_4_^+^ measurements. This was also the case for most major cations and trace elements measured (Table S2). Nitrate was found to be lower in the summed-up fractions than in the bulk soil samples, which is likely explained by leaching of NO_3_^-^ during fractionation, due to its high aqueous solubility. The observed median diameters of the PSFs were within the expected size range, except in the < 2 μm fraction where d_50_ was 3.1.

**Table 1:**
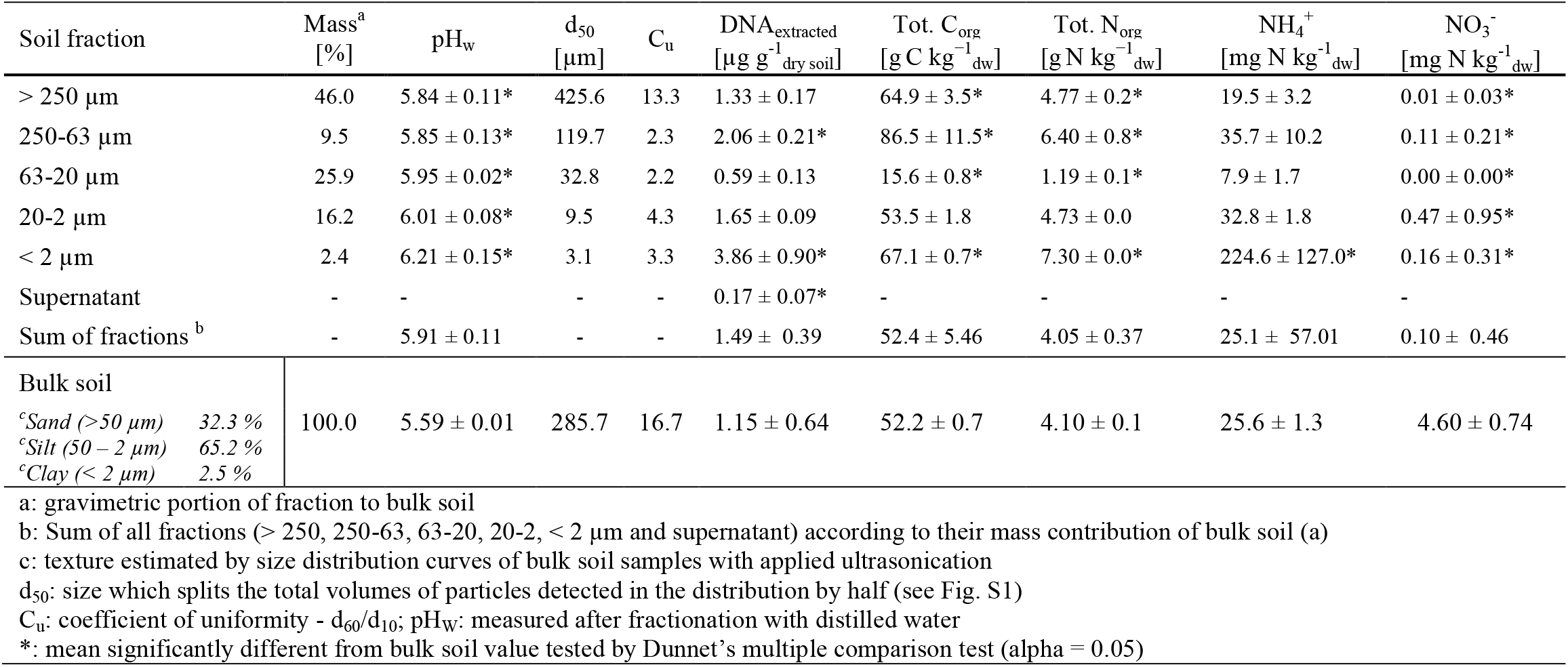
Summary of main soil physical and chemical parameters as well as extracted DNA of PSFs and bulk samples of Rothamsted Park Grass soil.

### Microbial community structures in PSFs

NMDS plots of RISA band profiles from PSFs and bulk soil samples (Fig. 2) showed that bacterial as well as fungal community structures differed between the sample groups. Permanova tests and subsequent multivariate homogeneity of group dispersions analysis on Bray-Curtis dissimilarity matrices (Table S3) showed that the group means were significantly different in the data set. However, it could not be excluded that these differences were due to a heterogeneity of variance among the different groups.

**Fig. 2:**
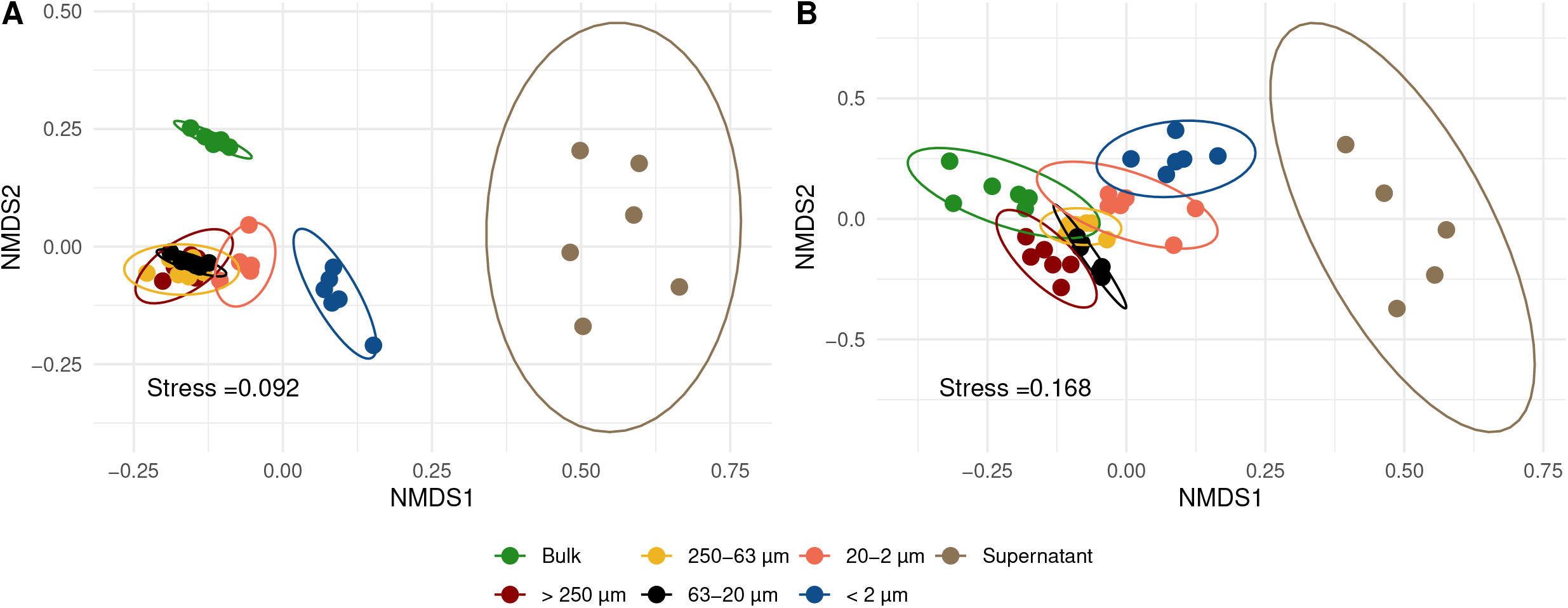
Non-metric multidimensional scaling (NMDS) of bacterial (A) and fungal (B) RISA band profiles of different PSFs and bulk soil samples based on Bray-Curtis dissimilarity matrices. Each point represents one replicate (n=6) and confidence ellipses are plotted for each sample group.

Results from annotated bacterial and fungal SSU rRNA amplicons are shown in Fig. 3 and 4 respectively. The order of taxa on the y-axis as well as the PSFs and bulk soil samples on the x-axis of the heatmap plots are based on a PCoA ordination of unweighted unifrac distances and therefore the left and right edges of the plots are adjacent in a continuous ordering and more discriminating groups are placed in the middle of the heatmap. Taking all bacterial ASVs at genus level into account (Fig. 3, panel A), PSFs with bigger aggregated structures (> 250 and 250-63 μm) showed communities closer to bulk soil samples based on relative abundance. The smaller PSFs (< 63 μm) showed a distinct community composition which placed them central of the heatmap plot. The supernatant clearly separated from all other PSFs with only a few taxa dominating the community. Here, almost all bacterial phyla detected in the PSFs decreased in relative abundance in the supernatant fraction except for Bacteroidota (mainly *Mucilaginibacter* and *Edaphobaculum*) and Proteobacteria (mainly *Pseudomonas*, *Massilia*, *Duganella*, *Janthinobacterium* and *Variovorax*) (Fig. 3, panel D). The relative abundance of *Pseudoathrobacter* (Actinobacteriota) was also increased in the supernatant and the < 2 μm fraction. When we analyzed the dataset only taking higher and lower abundant (> 1 and < 1 % relative abundance; Fig. 3, panel B and C, respectively) taxa into account, we saw similar patterns in the sample grouping on the x-axis.

**Fig. 3:**
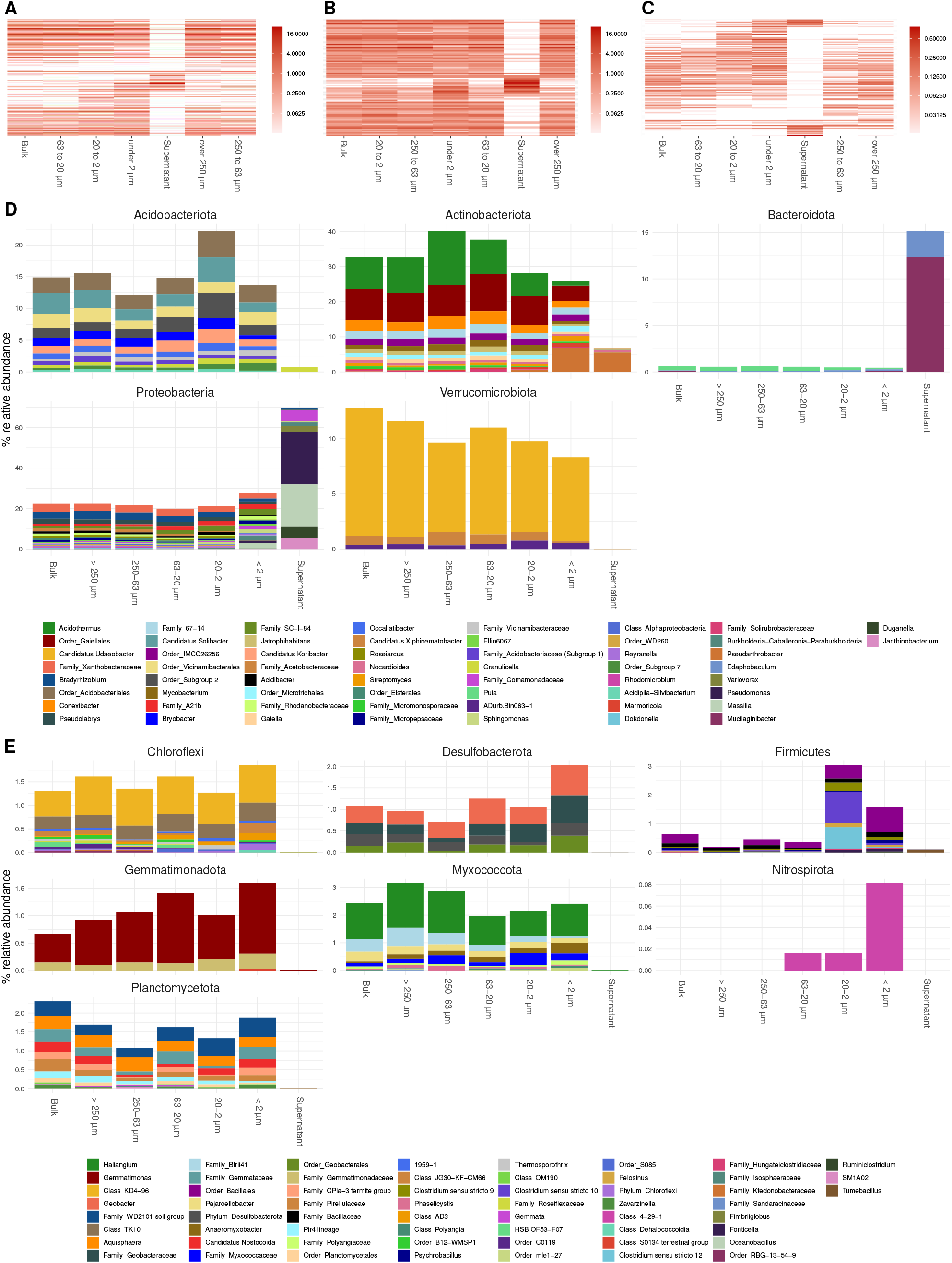
Annotated bacterial ASVs of SSU rRNA gene amplicons collapsed to genus level. A, B and C show heatmaps of all genera detected, genera present over and under 1 % in relative abundance, respectively. Order of x and y-axis of heatmaps are based on a PCoA ordination of unweighted unifrac distances. D: bar plots of higher abundant phyla showing genera over 0.3 % in relative abundance. E: bar plots of lower abundant phyla (unfiltered). If taxa unit could not be annotated at the genus level the name of the last higher rank where an annotation at a bootstrap value of 0.8 was possible is shown.

**Fig. 4:**
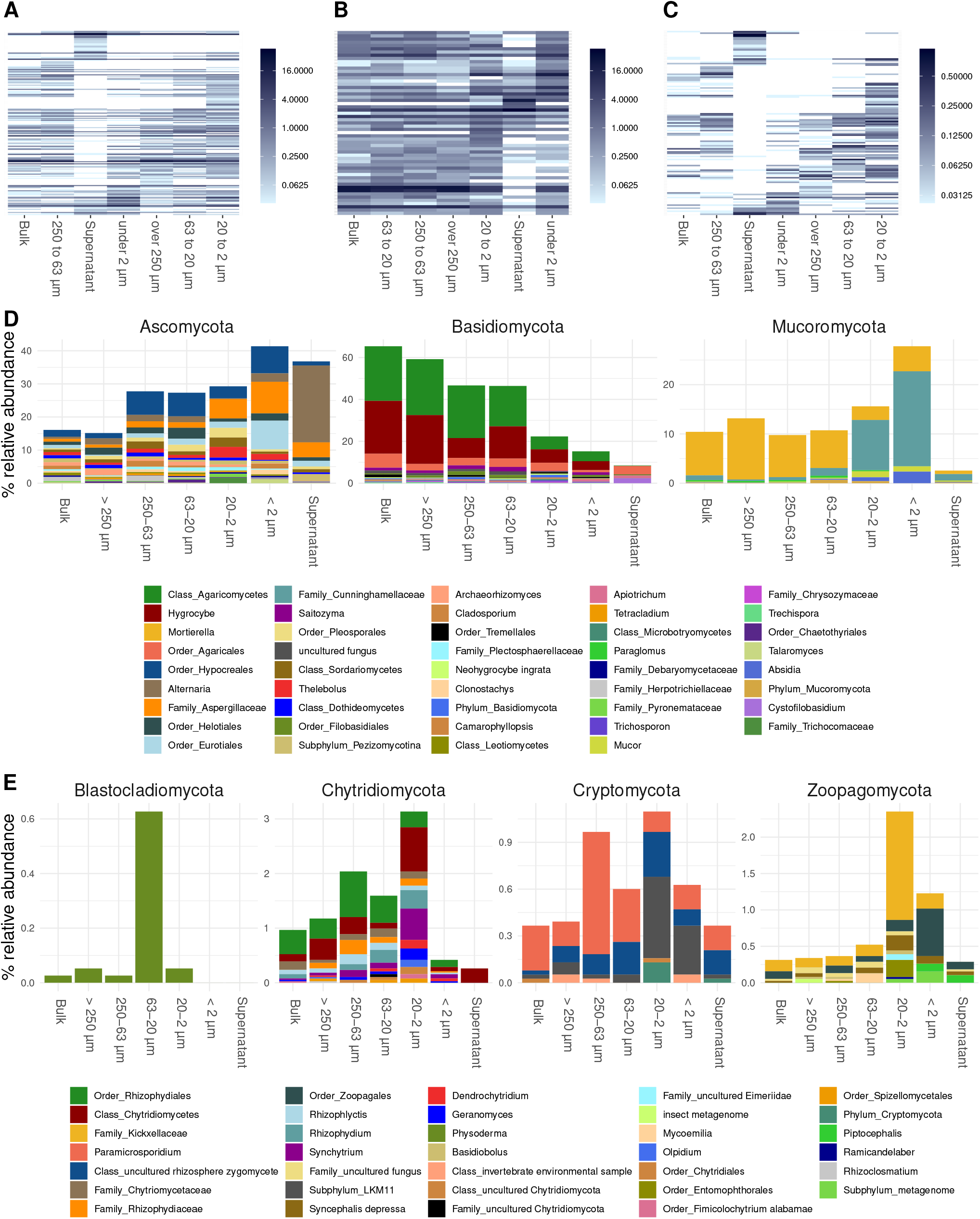
Annotated fungal ASVs of SSU rRNA gene amplicons collapsed to genus level. A, B and C show heatmaps of all genera detected, genera present over and under 1 % in relative abundance, respectively. Order of x and y-axis of heatmaps are based on a PCoA ordination of unweighted unifrac distances. D: bar plots of higher abundant phyla showing genera over 0.2 % in relative abundance. E: bar plots of lower abundant phyla (unfiltered). If taxa unit could not be annotated at the genus level the name of the last higher rank where an annotation at a bootstrap value of 0.8 was possible is shown.

Fungal communities showed a shift in community structure with particle size from larger to smaller PSFs at the phylum level (Fig. 3, panel D). The relative abundance of Basidiomycota (mainly Hygrocybe and other Agaricomycetes) decreased by about 45 % (from > 250 to < 2 μm), whereas that of Ascomycota (mainly Hypocreales, Eurotiales, Aspergillaceae and Alternaria) and Mucoromycota (mainly Mortierella and Cunninghamellaceae) increased within these size classes by about 25 and 15 %, respectively. Zoopagomycota were detected at higher levels in PSFs below 20 μm, albeit at overall low abundances (~ 0.1-1.5 % rel. abundance). Within the Mucoromycota annotated reads, a shift from Mortierella to taxa in the Mucorales order (Cunninghamellaceae) was observed in smaller PSFs. Some ASVs in the < 2 μm PSF were annotated to the genus Absidia. All fungal phyla showed a decrease in relative abundance in the supernatant samples with the exception of the Ascomycota, which were dominated by Alternaria and Aspergillaceae reads. Heatmap plots did not show a clear discriminating pattern between bigger and smaller aggregated structures. However, smaller PSFs were often placed next to the supernatant fraction which discriminated the most from bulk soil samples.

### Abundance of bacterial, fungal and N-cycling genes

The highest abundances of bacterial and fungal *SSU rRNA* gene copies per gram of dry soil estimated by qPCR were found in the < 2 μm (Fig. 5, panel A and B) fraction. The lowest were found in the 63 - 20 μm and in the supernatant. The other fractions showed similar abundances to the bulk soil samples. The proportion of nitrogen cycle marker genes in reductive pathways (nitrite reductases (*nirK* and *nirS*) and nitrous oxide reductases (*nosZ* clade 1 and 2; Fig. 2 panel C and D, respectively) relative to the total estimated abundance of bacteria was higher in the > 250 μm and 250 - 63 μm PSFs compared to the bulk soil samples. Overall, the proportion of marker genes for reductive nitrogen cycling decreased with particle size (Fig. 5 panel G; R^2^= 0.46). Marker genes for oxidative pathways in the nitrogen cycle (bacterial and archaeal ammonia monooxygenase (*amoA*); Fig. 5 panel E and F) did not change across the distinct sample groups and were generally low (~10^4^ copies g^-1^_dry soil_). However, we were not able to detect ammonia oxidizing bacterial genes in the < 2 μm and supernatant fractions, whereas this was not the case for the ammonia-oxidizing archaeal genes.

**Fig. 5:**
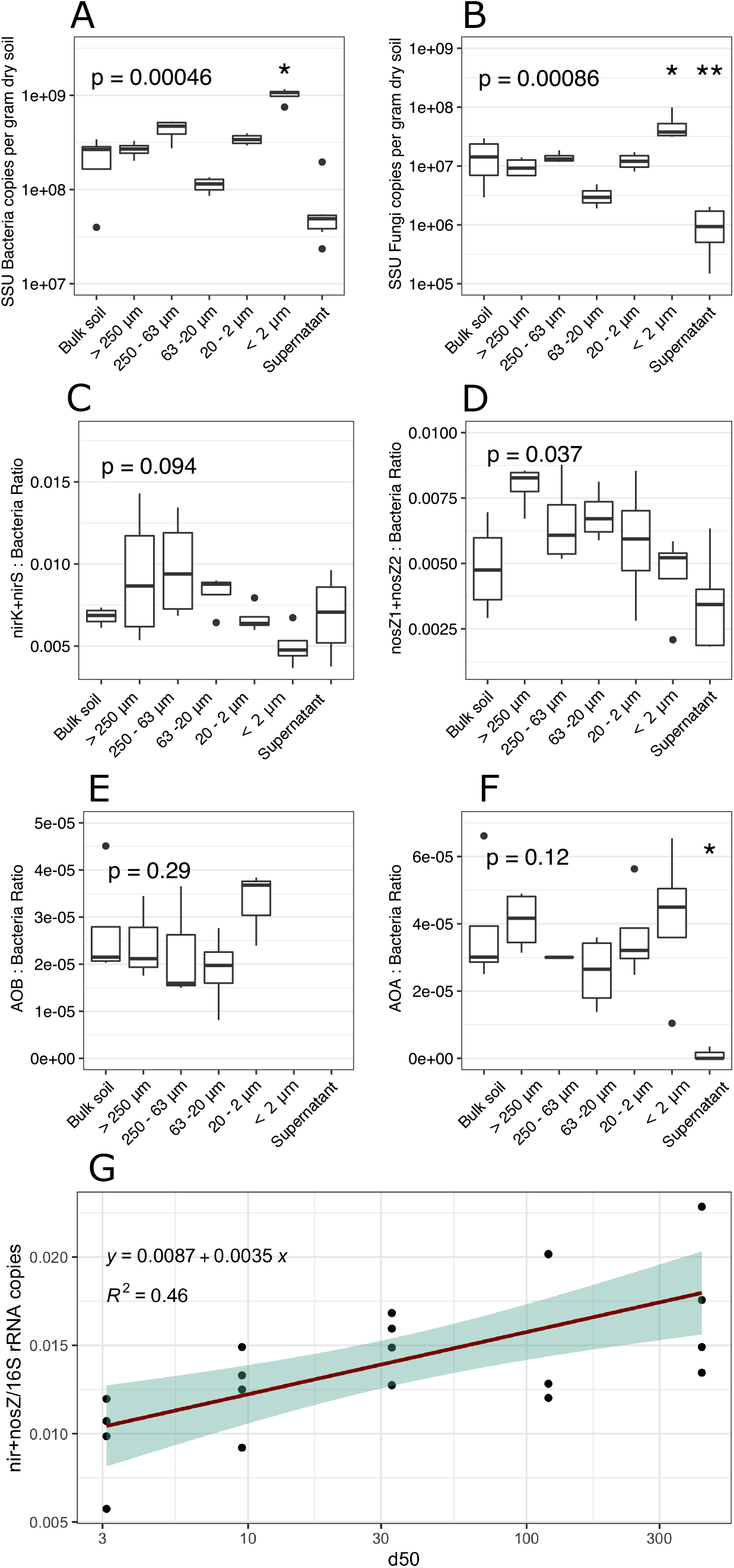
Abundance estimated by quantitative PCR (n=6) of bacterial and fungal SSU rRNA genes (A and B respectively) and ratios of various gene abundances in obtained fractions and bulk soil: C: nitrite reductases to bacterial SSU rRNA; D: nitrous oxide reductases to bacterial SSU rRNA; E: ammonia oxidizing bacteria to bacterial SSU rRNA; F: ammonia oxidizing archaea to bacterial SSU rRNA; G: linear regression of ratio of summed up reductive N-cycle marker to bacterial SSU rRNA. Values presented in the supernatant group were obtained by calculating the amount washed-off DNA from one gram of soil during the fractionation procedure to allow comparison. P-values correspond to non-parametric ANOVA Kruskal Wallis tests; asteriks indicate significance from Wilcoxon tests between the sample group and bulk soil samples (* and ** stand for alpha = 0.05 and 0.01 respectively).

## Discussion

Soil is one of the most diverse and physically complex microbial habitats on the planet (2). Aggregate formation from soil particles has generally been studied at the macroaggregate level (>250 μm) and has been shown to be a hierarchical process involving a series of biophysical processes (7, 21, 28). In these studies, aggregate size preferences and stabilization capacity have been shown to vary among fungal taxa and are related to specific physiological traits such as mycelial/hyphal density and enzyme and polysaccharide production (19, 27). In the microaggregate range, less is known, although these size classes are more relevant to the individual organisms themselves.

### Quality of soil fractionation

The perfect fractionation of a soil sample would result in each parameter measured in the fractions adding up to the value found in the bulk soil when multiplied with its mass fraction and totaled. Calculations in Table 1 and S2 show that this was the case for almost all measured parameters. However, if the totaled value was found higher or lower than the one in the bulk soil, this could indicate a cross-contamination between fractions if the measured analyte would be transferred to a fraction with a high (i.e., > 250 or 63-20 μm) or a low mass fraction like the < 2 μm one, respectively. For extracted DNA, the totaled value was found higher but not significantly different from the bulk soil sample. High amounts of extracted DNA and abundance of bacterial and fungal communities was found in the < 2 μm fraction. If these high levels would be caused by a significant transfer of microbial biomass or DNA to the < 2 μm fraction, we would expect a lower totaled value than the one obtained in the bulk soil sample due to the low mass fraction of the < 2 μm PSF (~2.5 %). This suggests that DNA based parameters are connected to the respective fractions in this study and are not critically biased by cross-contamination effects. It should be mentioned that this estimation of fractionation quality still has to be handled with care since multiple cross-contamination between fractions could cancel each other out. However, since these effects are difficult to trace back in these kinds of studies, we believe that this approach adds a level of confidence to our dataset.

### Microbial size preferences at the microscale level

We examined microbial community composition within soil particle size fractions, with a focus on microaggregates. Based on our data, we observe a change in fungal community structure across the different PSFs. A change was also observed for bacteria, but only in microaggregates smaller than 20 μm and in the supernatant. This suggests that fungi play an important role in aggregate formation and soil structuring, which is supported by a recent study showing that fungi are more crucial for soil structuration than bacteria at the microscale (i.e. smaller than 53 μm), and suggesting that fungi are the most important agents of selforganization (29). However, other studies concluded that fungi are more associated to macroaggregates because of their hyphae (30, 31), but only a few studies have specifically examined fungi in microaggregates.

#### Over 250 and 250 - 63 μm

The shift to smaller particle sizes induced by ultrasonic (US) treatment showed that we were able to collected stable aggregates in these PSFs (Fig. S1). The overall soil structure calculated after US dispersion (32.3, 65.2 and 2.5 % for sand, silt and clay respectively; Table 1) is still shifted to larger particles compared to the conventional soil structure analysis where chemical and mechanical dispersion of each soil particle is approximated (19, 58 and 23 % for sand, silt and clay, respectively (32)). This confirms that the strong bonds between particles remained intact during our procedure and qualifies our samples for microbiologically meaningful interpretation. Aggregate stability has been linked to hyphal abundance of saprophytic and lignin degrading basidiomycetes previously (26, 27). Basidiomycota *(Hygrocybe* and likely close relatives within the Agaricomycetes) dominated the fractions larger than 250 and between 250 - 63 μm but decreased in smaller PSFs. *Hygrocybe* was found before in nutrient poor, unmanaged grasslands (33–35). Members of the Agaricales are known to exhibit a mycorrhizal or biotrophic lifestyle (9, 36, 37). These fungi can establish large and long lived mycelial networks, which likely have a stabilizing effect in the aggregates obtained in these PSFs through hyphal meshing of smaller microaggregates and production of binding agents (38–40).

Ascomycota were observed to increase in relative abundance with decreasing particle size (Fig. 4; panel D). This phylum has been shown to be particularly efficient at forming aggregates at a macroscopic scale, due to its high hyphal density (19). However, our data suggests that small microaggregates constitute preferred habitats for Ascomycota were specific taxa increased in relative abundance like Aspergillaceae and Eurotiales.

#### 63 - 20 μm

This size class was characterized by organomineral silt microaggregates that had the lowest concentrations of organic carbon and the second highest mass fraction of 25 %, but the lowest values of measured ions (Table S2), extracted DNA, and gene copy number. These data suggest the presence of inert particles with a low negative surface charge that would limit adhesion of organics, ions and microbes (41, 42). Blastocladiomycota, although found only in low relative abundance (~ 0.02 %) in bulk soil communities, were preferentially detected in these microaggregates. These organisms, along with Chytridiomycota, belong to the zoosporic true fungi group (43). Their ability to produce zoospores with a flagellum (44, 45), their survival in nutrient depleted environments for long periods of time (46) and their attachment to mineral surfaces and insoluble minerals may explain why they are mainly detected in the silt-organomineral microaggregates.

#### 20 - 2 μm

Strikingly, a difference in bacterial community composition was observed in PSFs below 20 μm which corresponds to the distance where the majority of bacterial interactions occur in soil (Fig. 2 and 3) (20). Chytridomycota and Zoopagomycota were preferentially selected in this size class, albeit at low levels. Microaggregates in this size range are characterized by low amounts of new and readily available carbon, as they contain little or no plant debris (47). The cores of these small microaggregates are either organic debris (e.g. 46, 47) or bacterial cells (41, 50, 51) that are then surrounded by phyllosilicates and metal oxides (7, 10, 12). The quality of organic matter has been shown to decrease with aggregate size, as indicated by a decrease of the C:N ratio (7, 52–55), which is consistent with our data. Therefore, the ability of Chytridiomycota to degrade recalcitrant matter and withstand extreme environments could explain its presence in this fraction. Also, Chytridiomycota are rarely observed in soils, with the exception of unvegetated soils at high altitude (56) where they have been found to be dominant. Zoopagomycota, on the other hand, are small fungi characterized by pathogenic or symbiotic associations with animals and fungi with virtually no association to living plants (57). Their presence in aggregates may be related to their small size and pathogenic lifestyle. Another explanation for the accumulation of Zoopagomycota could be that, besides the flagellated short lived spore forms, many members in this group can form long-term resting spores (~10 - 20 μm in diameter) which are possibly collected in this fraction (57). Overall, the observed distribution of rare fungal phyla within microaggregate fractions suggests that the applied method could potentially be used to study these organisms in soil.

A shift within the Mucoromycota was observed from Mortiella dominating bigger PSF to Mucorales related taxa (Cunninghamellaceae, Mucor and Absidia) below 20 μm. Mucoraceae have previously been reported to dominate non-Dikarya in this fraction and have recently been found to establish symbiotic relationships with plants (58, 59). However, the increase of these fungi in small microaggregates remains speculative. Following our observation of a general decrease of Basidiomycota involved in enmeshment of larger aggregate classes, different adhesive forces might take over in order to connect single particles to microaggregates, such as excretion of exopolysaccharides (EPS), for attachment (6, 22, 27, 60, 61). A possible mechanism involving the binding of the polymer chitosan present on cell walls of Mucorales related strains to clay particles is described in the < 2 μm section below.

A change in bacterial community was also observed in the 20-2 μm fraction, with an increased abundance of *Clostridium* sequences for example. This could indicate oxygen-limited conditions, as *Clostridium* was previously linked to the fermentation of organic matter in the soil (62). However, as *Clostridium* is spore forming, its elevated abundance could also be due to an accumulation of spores in that fraction. Bacterial taxa have been shown to exhibit habitat preferences within and at the interfaces of microaggregates (63–69), but these preferences have generally not been detected for the dominant soil-colonizing taxa, but rather for some low-abundance taxa (65).

#### Below 2 μm

Bacterial and fungal community structures changed in this PSFs compared to the larger ones together with a significantly higher abundance of microbes, accumulation of organic carbon and nitrogen and important cationic nutrients such as ammonium, iron, calcium, copper and chromium. Several studies have also found that clay fractions contained the highest gene copy numbers or biomass (13, 66, 70). While the high numbers of fungi could be explained by an accumulation of spores in this fraction, as suggested by Hemkemeyer et al. 2019 (58), we were surprised to find that the relative abundance of Mucoromycota increased to ~27 %. The cell wall of Mucorales increasing in this PSF is considered unique as it is the only natural source of the biopolymer chitosan known to date (in contrast to chitin, which is abundant) (71–74). Chitosan has been intensively studied in engineering applications due to its binding properties to clay, resulting in nanocomposites with unique properties such as strong absorption (75). It seems plausible that a similar mechanism is responsible for the increased binding of Mucorales to clay particles in soil, which would explain the observed accumulation in this size class. In general, clay particles have varying degrees of negative charges on their surface depending on their chemical composition and are known to interact with microbes in multiple ways (42, 76). This allows microbial cells to adhere through electrostatic forces where enhanced adhesion through shedding of EPS-like structures could occur under the right conditions (41, 60). EPS exhibit and enhance key criteria for microbial life such as nutrients and water binding, physical and chemical protection and signaling (6, 11). It is therefore not surprising that microbes thrive in close proximity to each other, which may explain the high abundance of bacteria and fungi found in the < 2 μm fraction. Also, the observed changes in taxonomical composition of bacteria and fungi in this fraction indicates unique niches potentially created by biofilm like community assembly/dependency.

#### Supernatant

Almost all of the taxa separated by centrifugation of the < 2 μm fraction were previously reported to be involved in EPS production and biofilm formation (22, 61, 77–82) and dominated by the phyla Proteobacteria (*Pseudomonas*, *Massilia*, *Variovorax*, *Burkholderia, Duganella and Janthinobacterium*) and Bacteroidota (*Mucilaginibacter* and *Edaphobaculum*) which were present in the PSFs only at low abundances (Fig. 3 and S2). Adding centrifugation steps to soil fractionation protocols could therefore be of use in studies focusing on isolating biofilm forming microorganisms.

The settlement of particles in a solution depends in part on density (83, 84). Thus, one explanation for the observed occurrence of specific taxa could be the presence of gas vesicles, which can provide buoyancy (85, 86). Extracellular DNA (eDNA) could also remain in solution rather than settling during centrifugation. In a recent study, it was shown that *Pseudomonas* forming biofilm on fungal hyphae used eDNA filaments to create stability (87). This could explain the higher abundance of *Pseudomonas* sequences in the supernatant, although it would imply that the excreted DNA contains the quantified and sequenced rRNA gene. However, the amount of 16S copies per ng DNA were found to be similar in the supernatant and bulk soil samples, suggesting a comparable ratio of rRNA gene to total DNA.

Filamentous fungi were not expected to be detected in the supernatant, or if so only in low abundances. Indeed, the abundance of fungi in this fraction was found to be significantly lower (Fig. 5, panel B) compared to the PSFs. The dominance of Alternatia related sequences is likely linked to spores from this genus which do not settle during the final centrifugation step. Yeasts or fungi which can reproduce in yeast form might also end up in this fraction, albeit at lower abundances. This was the case for many ASVs only found in the supernatant fraction which were annotated to yeasts including *Cystobasidium*, *Malassezia*, *Sporidiobulus*, *Schefferomyces, Pichia* and *Filobasidium* (Figure S3, panel B). *Saitozyma*, another ubiquitous yeast often found in soil (88), was linked to bigger PSFs (Figure S3, panel A) which could point to differences in preferred habitats and consequently attachment forces of yeast in soil.

### Nitrogen cycling potentials in microhabitats

Given the existence of heterogeneous niches at the microscopic scale and the low oxygen levels found in stable aggregates (5), we hypothesized that the denitrification potential would vary among the different size fractions. The proportion of nitrite reductase (*nirK* and *nirS*) and nitrous oxide reductase (*nosZ* clade 1 and 2) copies to the overall bacterial community was consistent with previous studies on unmanaged soils (89–93). The observed relationship of a potential N-oxide reducing community decreasing in relative abundance with particle size (Fig. 5, panel G) suggests that denitrifying organisms find optimal conditions in larger stable aggregate structures. This is in agreement with a previous study, which showed that denitrification activity was higher in larger PSFs while copy number of marker genes was comparable (94). In contrast, nitrogen cycle oxidative pathways are uniformly distributed in PSFs, albeit at low levels. The low abundance of ammonia oxidation has been previously demonstrated in pristine grasslands compared to managed soils (95–97). In unfertilized soils, nitrogen mineralization from decomposing organic matter and biological N-fixation are critical for N-supply (98, 99). Ammonium has been found to accumulate in the < 2 μm fraction, which is likely the result of a high cation exchange capacity of clay and potential EPSs (60, 100). However, the low abundance of amoA genes detected suggests that this NH_4_^+^ pool was not accessible to ammonia oxidizers and thus could be primarily used in an assimilatory manner by plants and other microbes (101).

### Conclusions

<u>Our data confirm that distinct habitat preferences of microbial communities are reflected by the size of soil particle aggregates and extend this by showing that a further separation of microaggregates (< 250 μm) leads to bigger observed discrimination of community structures. This shows potential for soil studies looking at scales of a few micrometers as this might better reflect the functional relationships of communities than bigger structures. Advances in data generation from smaller samples in combination with sampling techniques at micrometer scale will therefore be key to knowledge gain in a complex environment as soil</u>.

The observed shift in fungal taxa confirms the proposed role of Basidiomycota in stabilizing larger aggregates from previous studies. However, the changes in fungal communities observed below 20 μm point to a different role of fungi at those scales which potentially can be answered in bacterial-fungal interactions as these are often found to be dependent on close proximity (102–104). Furthermore, the selection for fungal phyla usually found at low abundances in various PSFs point to the potential of similar fractionation approaches to study these organisms.

As bacteria are present as single cells or small colonies they are not involved in the physical stabilization of bigger aggregates as fungi are by hyphal enmeshment which was confirmed by the overall similar community structures found in particles above 63 μm. However, below 20 μm changes were observed indicating the access to habitats with more specific conditions than in bigger structures. Surprisingly, only a few taxa from two bacterial and one fungal phyla (Proteobacteria, Bacteroidota and Ascomycota, respectively) were recovered in the supernatant fraction, while others (i.e. Acidobacteria and Verrucomicrobia) were almost exclusively found in PSFs. As we rinse soil through sieves during the applied wet fractionation, these microbes are likely washed-off the soil matrix and thus can be seen as loosely attached. Therefore, we believe that the probability of cross-contamination in these kinds of studies has to be evaluated differently for each obtained fraction. While loosely attached microbes in the supernatant likely stem from particles over the entire size range, communities found in the bigger PSFs can be linked to stable cores of bigger aggregated structures. The biggest risk of cross-contamination effects might be in the very fine < 2 μm fraction. Microbes and extracellular DNA might bind to dispersed clay particles until they are saturated during the fractionation, which could lead to misinterpretation (76, 105). The usually high microbial densities and diversities found in that fraction would support such a scenario in wet fractionation approaches (58, 70). Strikingly, in our study, taxa found in the supernatant fraction were often also detected in elevated relative abundances in the < 2 μm PSF. It seems plausible that this is caused by an efficient detachment of i.e. *Pseudomonas, Massilia* or *Pseudoathrobacter* from the soil surface followed by binding to clay particles until saturation and the remaining DNA is found in the supernatant. However, data from Bacteroidota and fungi do not support this theory. It has previously been shown that the nature of the fractionation method has an influence on the distribution of extracellular enzyme activity, which is likely also the case in studies focusing on microbes in soil (16). As dry fractionation approaches usually obtain no clay fractions, a comparison of data to estimate these cross-contamination effects proves difficult.

Overall, the fractionation of soil in micro-fractions shows potential to answer relevant research questions in soil microbial ecology by accessing communities at scales where they operate. Specifically, the attempt to wash-off and analyze microbes with varying degrees of detachment forces from soil might harbor potential to gain insight in how communities are organized in biofilm-like structures in soil.

## Material and Methods

### Soil sampling and physical fractionation

Samples were collected from the untreated control plots of the Park Grass Experiment, Rothamsted Research, Hertfordshire, UK (35) in October 2013. These control plots have not been manipulated (i.e. tillage, fertilization and liming) for over 150 years at the time of sampling. Therefore, the microbial communities and their spatial distribution at the microscale can be interpreted as a results of natural soil structuring processes. Soil samples were fractionated in PSFs and analyzed in parallel with a non-fractioned sample (referred to as “bulk soil” throughout the manuscript). For fractionation, the soil was sieved (2mm) and 6 PSFs were obtained using a procedure adapted from Jocteur Monrozier et al., 1991 (13) to optimize preservation of soil microaggregation (beads were not used in the macroaggregate disruption step). Autoclaved and sterilized material and solutions were used to avoid contamination of samples. Ultra-pure water (200 mL) was added to 8 replicates (6 for biological, 2 for physical and chemical analysis) of 30 g of bulk soil in 500 mL Duran^®^ bottles and subsequently placed on an orbital shaker for 1h at 250 rpm. This gentle soil dispersion was repeated 3 times. Subsequent fractionation steps were carried out as previously described and consisted of a sequence of i) wet sieving of the dispersed soils to separate fractions of particle sizes > 250 μm and 250-63 μm, ii) settling of 63-20 μm diameter particles at 1 g (approx. 10 min), iii) sedimentation of 20-2 μm diameter particles at 1g (5 to 6h) and iv) centrifugation at 11,000 g to collect particles of < 2 μm in diameter (13, 106). The supernatant from the centrifugation step was included in the further biological analysis and represented all organisms washed from the soil matrix during the procedure and not settled during centrifugation. All fractions were frozen at −80 °C for subsequent DNA extraction and chemical analysis. Soil physical analysis was carried out directly on fractions after the completed fractionation procedure.

### Soil physical and chemical analysis

The particle size distributions of bulk soil and solid fractions were determined by laser granulometry with a Mastersizer 2000 laser granulometer (Malvern Instruments Ltd, United Kingdom). A refractive index of 1.55 was used as a model parameter according to the procedure described by Navel et al. 2014. Total organic C and N concentrations of bulk soil and PSFs were determined in duplicate on approximately 15 mg of blended soil samples with a FlashEA1112/FLASH 2000 Elemental Analyzer equipped with a thermal conductivity detector. For ammonium (NH_4_^+^) and nitrate (NO_3_^-^) measurements, 1 g dry weight of bulk soil or PSF was extracted with 1 M KCL solution (1:5, w:v) for 30 min on a spinning wheel, subsequently centrifuged (5 min at maximum speed) and the supernatant used for quantification. Colorimetric measurements were performed as described previously (107). Major and trace elements were quantified in duplicate with 0.2 g of air-dried bulk soil and PSFs samples after microwave-assisted acid digestion. Samples were suspended in a mixed solution of HF:HNO_3_ (6:2, v:v) in a pressurized closed-vessel microwave system (NovaWAVE, SCP SCIENCE). The digestion program consisted of a 10-min gradual increase to 180°C, a 10-min step at 180°C and then a cooling step. Mineral elements were quantified by ICP-AES (Perkin Elmer; Optima 3000 DV).

### DNA extraction and sequencing

Six replicates of homogenized bulk soil and PSF samples (100 - 300 mg dry soil) were used for DNA extraction (DNeasy Power Soil extraction kit, QIAGEN). The water content of each sample was determined gravimetrically by drying 1 g of sample at 60°C for 24 h. The supernatant samples were filtered on sterile cellulose membranes (MF-Millipore™ membrane filters; pore size 0.45 μm) with a sterile filtration unit and DNA was subsequently extracted from the membranes (DNeasy PowerWater Kit, QIAGEN). Isolated DNA was quantified with the Qubit^®^ dsDNA HS Assay Kit on a Qubit^®^ Fluorometer (Invitrogen™) and stored at −20°C for further analysis.

### RISA analysis

Internal transcribed spacer regions (ITS) of bacterial and fungal communities were amplified from isolated DNA with primer pairs FGPL 132-38 / FGPS 1490-72 and fITS7 / ITS4 respectively (108, 109). Each 25 μL reaction contained 12.5 pmol of the primers and 2 ng of template DNA. Cycling conditions for the bacterial assay were: 94°C for 10 min; 29 cycles of 94 °C for 20 s, 55 °C for 30 s and 72 °C for 1.5 min; final elongation step at 72 °C for 10 min. Cycling conditions for the fungal assay were: 94°C for 10 min, 35 cycles at 94 °C for 20 s, 57 °C for 30 s and 72 °C for 1.5 min; final elongation step at 72 °C for 10 min. PCR products were run on DNA 1000 chips on an Agilent 2100 Bioanalyzer system (Agilent Technologies, Inc.). Tables of band profile traces were imported in R and NDMS on Bray-Curtis dissimilarity matrices were performed with the *metaMDS* function from the ‘vegan’ package and plots generated with the ggplot2 package (110). PERMANOVA analysis was done on Bray-Curtis dissimilarity matrices with the *adonis2* function with 999 permutations. Test on multivariate homogeneity of group dispersions was done by the *betadisper* and *permutest* function of the vegan package.

### Amplicon sequence generation and analysis

RISA band profiles suggested distinct community profiles in the fractions. DNA samples were pooled according to fractions before library preparation for amplicon sequencing as follows: Fungal V7-V8 region of the *18S rRNA* gene was amplified using FR1/FF390 primers (5’-AICCATTCAATCGGTAIT-3’ – 5’-CGATAACGAACGAGACCT-3’; 348 bp amplicon). Bacterial V4-V6 region of the *16S rRNA* gene was amplified using 16S-0515F / 16S-1061R primers (5’-TGYCAGCMGCCGCGGTA-3’ -- 5’-TCACGRCACGAGCTGACG-3’; 540 bp amplicon). Primers were tailed with Roche multiplex identifiers (MIDs). Each reaction mix (50 μL) contained 10 pmol of each primer, 40 nmol of dNTPs, 1 x reaction buffer, 100 ng of template DNA and 1 x Titanium Taq DNA Polymerase (Invitrogen™). Cycling conditions were: 95°C for 1 min; 35 cycles of 95 °C for 30 sec and 68 °C for 1 min; final elongation step at 68 °C for 3 min. PCR reactions were loaded on an agarose gel and bands cut out and purified (Gfx – DNA Purification Kit, GE-Healthcare). Concentration of DNA was quantified with the Qubit^®^ dsDNA HS Assay Kit on a Qubit^®^ Fluorometer (Invitrogen™). An equimolar pool of amplicon samples (60 ng DNA of each sample) was sent for 454-pyrosequencing (Beckman Coulter Genomics).

All steps conducted from raw sequence files to phyloseq objects used to create plots are documented step by step in the supplemental material. Briefly, initial quality filtering and sequence primer trimming was performed with PRINSEQ and pyrosequencing pipeline of RDP (111, 112). Further steps followed standard analysis steps in the dada2 pipeline using recommended parameters for reads generated with 454-pyrosequencing. For bacterial amplicons, 187465 raw reads gave 116090 filtered, 103405 denoised and 99814 chimera filtered reads which resulted in 717 amplicon sequence variants (ASVs). For fungal 18S rRNA gene amplicons, 135025 raw reads gave 130136 filtered, 126333 denoised and 91080 chimera filtered reads which resulted in 1216 ASVs. The training set for taxonomic annotation of prokaryotic reads was downloaded from the dada2 repository. For fungal 18S annotation, a training set was generated from the SILVA SSU reference database cluster to 99% similarity including only fungal reads and taxonomic strings in the fasta-header were converted to be compatible with the dada2 annotation algorithm. Detailed steps involved in database generation are documented in the supplemental material. Phylogenetic trees used in the analysis were generated with FastTree from multiple sequence alignments of ASV sequences generated with MAFFT (113, 114).

### Quantitative PCR

Standard curves for all reactions were derived from serial dilutions of linearized pGEM-T plasmids with the target inserted sequences (90). All standard curves were linear and had comparable R-square (0.96 to 0.99) and efficiency (0.85 to 1.05) values. Quantitative PCR was performed on a Corbett Rotor-Gene 6000 real-time PCR cycler and QuantiTect SYBR^®^ Green PCR Kit (Quiagen). Each reaction (25 μL) contained 12.5 μL 2x QuantiTect SYBR^®^ Green Mix; 0.3 to 1.8 μL of each primer (10 μM); 100 ng of T4 gene protein 32 (Thermo Fisher Scientific Inc.); 2 to 5 ng of template DNA and PCR grade water. Primer sequences are summarized in Table S1. In each run 3 non-template control reactions were included to test for contamination of the master mix. Melting curve analysis was used as quality control of qPCR runs. Results from runs showing no amplification in the non-template controls and only one melting peak overlapping with the peaks from standards were considered for further analysis. Plotting and statistical analysis were performed in R (110).

## Acknowledgements

The authors acknowledge the help of Frédéric Lehembre in experimental design and soil chemical analysis. We would also like to thank Laurent Philippot and David Bru from the Department of Agroecology in Dijon for fruitful discussion of the results and provision of qPCR standards used in this study, respectively. We would also like to thank Sandrine Demaneche and Lorna Dawson for access to samples from the Rothamsted Park Grass experiment. Chemical analytical aspects were supported at IGE by the Air-O-Sol and MOME technical platforms, within the Labex OSUG@2020 (ANR10 LABX56).

## Author contributions

Conceptualization: C.K., C.L., P.S. and J.M.; Analysis: C.K. and A.N.; writing – original draft: C.K.; writing – review & editing: all authors.

## References

1. Bardgett RD, Van Der Putten WH. 2014. Belowground biodiversity and ecosystem functioning. Nature 515:505–511.

2. Young IM, Crawford JW. 2004. Interactions and Self-Organization in the Soil-Microbe Complex. Science (80-) 304:1634–1637.

3. Rabot E, Wiesmeier M, Schlüter S, Vogel HJ. 2018. Soil structure as an indicator of soil functions: A review. Geoderma 314:122–137.

4. Ebrahimi A, Or D. 2016. Microbial community dynamics in soil aggregates shape biogeochemical gas fluxes from soil profiles - upscaling an aggregate biophysical model. Glob Chang Biol 22:3141–3156.

5. Sexstone AJ, Revsbech NP, Parkin TB, Tiedje JM. 1985. Direct Measurement of Oxygen Profiles and Denitrification Rates in Soil Aggregates. Soil Sci Soc Am J 49:645–651.

6. Costa OYA, Raaijmakers JM, Kuramae EE. 2018. Microbial extracellular polymeric substances: Ecological function and impact on soil aggregation. Front Microbiol 9:1636.

7. Totsche KU, Amelung W, Gerzabek MH, Guggenberger G, Klumpp E, Knief C, Lehndorff E, Mikutta R, Peth S, Prechtel A, Ray N, Kögel-Knabner I. 2018. Microaggregates in soils. J Plant Nutr Soil Sci 181:104–136.

8. Bronick CJ, Lal R. 2005. Soil structure and management: A review. Geoderma 124:3–22.

9. Rillig MC, Mummey DL. 2006. Mycorrhizas and soil structure. New Phytol 171:41 – 53.

10. Lehndorff E, Rodionov A, Plümer L, Rottmann P, Spiering B, Dultz S, Amelung W. 2021. Spatial organization of soil microaggregates. Geoderma 386:114915.

11. Watteau F, Villemin G. 2018. Soil microstructures examined through transmission electron microscopy reveal soil-microorganisms interactions. Front Environ Sci 6:1–10.

12. Lehmann J, Kinyangi J, Solomon D. 2007. Organic matter stabilization in soil microaggregates: Implications from spatial heterogeneity of organic carbon contents and carbon forms. Biogeochemistry 85:45–57.

13. Jocteur Monrozier L, Ladd JN, Fitzpatrick RW, Foster RC, Rapauch M. 1991. Components and microbial biomass content of size fractions in soils of contrasting aggregation. Geoderma 50:37–62.

14. Hemkemeyer M, Christensen BT, Martens R, Tebbe CC. 2015. Soil particle size fractions harbour distinct microbial communities and differ in potential for microbial mineralisation of organic pollutants. Soil Biol Biochem 90:255–265.

15. Zhang Q, Zhou W, Liang G, Sun J, Wang X, He P. 2015. Distribution of soil nutrients, extracellular enzyme activities and microbial communities across particle-size fractions in a long-term fertilizer experiment. Appl Soil Ecol 94:59–71.

16. Bach EM, Hofmockel KS. 2014. Soil aggregate isolation method affects measures of intra-aggregate extracellular enzyme activity. Soil Biol Biochem 69:54–62.

17. Bailey VL, Fansler SJ, Stegen JC, McCue LA. 2013. Linking microbial community structure to β-glucosidic function in soil aggregates. ISME J 7:2044–2053.

18. Bailey VL, McCue LA, Fansler SJ, Boyanov MI, DeCarlo F, Kemner KM, Konopka A. 2013. Micrometer-scale physical structure and microbial composition of soil macroaggregates. Soil Biol Biochem 65:60–68.

19. Lehmann A, Zheng W, Ryo M, Soutschek K, Roy J, Rongstock R, Maaß S, Rillig MC. 2020. Fungal Traits Important for Soil Aggregation. Front Microbiol 10:1–13.

20. Raynaud X, Nunan N. 2014. Spatial ecology of bacteria at the microscale in soil. PLoS One 9:e87217.

21. Wilpiszeski RL, Aufrecht JA, Retterer ST, Sullivan MB, Graham DE, Pierce EM, Zablocki OD, Palumbo A V., Elias DA. 2019. Soil Aggregate Microbial Communities: Towards Understanding Microbiome Interactions at Biologically Relevant Scales. Appl Environ Microbiol 85:1–18.

22. Pandit A, Adholeya A, Cahill D, Brau L, Kochar M. 2020. Microbial biofilms in nature: unlocking their potential for agricultural applications. J Appl Microbiol 129:199–211.

23. Donot F, Fontana A, Baccou JC, Schorr-Galindo S. 2012. Microbial exopolysaccharides: Main examples of synthesis, excretion, genetics and extraction. Carbohydr Polym 87:951–962.

24. Chenu C, Cosentino D. 2011. Microbial regulation of soil structural dynamics, p. 37–70. In The Architecture and Biology of Soils: Life in Inner Space.

25. Beare MH, Hu S, Coleman DC, Hendrix PF. 1997. Influences of mycelial fungi on soil aggregation and organic matter storage in conventional and no-tillage soils. Appl Soil Ecol 5:211–219.

26. Caesar-TonThat TC, Cochran VL. 2000. Soil aggregate stabilization by a saprophytic lignin-decomposing basidiomycete fungus I. Microbiological aspects. Biol Fertil Soils 32:374–380.

27. Tisdall JM, Nelson SE, Wilkinson KG, Smith SE, McKenzie BM. 2012. Stabilisation of soil against wind erosion by six saprotrophic fungi. Soil Biol Biochem 50:134–141.

28. Six J, Bossuyt H, Degryze S, Denef K. 2004. A history of research on the link between (micro)aggregates, soil biotas, and soil organic matter dynamics. Soil Tillage Res 79:7–31.

29. Crawford JW, Deacon L, Grinev D, Harris JA, Ritz K, Singh BK, Young I. 2012. Microbial diversity affects self-organization of the soil - Microbe system with consequences for function. J R Soc Interface 9:1302–1310.

30. Kandeler E, Tscherko D, Bruce KD, Stemmer M, Hobbs PJ, Bardgett RD, Amelung W. 2000. Structure and function of the soil microbial community in microhabitats of a heavy metal polluted soil. Biol Fertil Soils 32:390–400.

31. Kihara J, Martius C, Bationo A, Thuita M, Lesueur D, Herrmann L, Amelung W, Vlek PLG. 2012. Soil aggregation and total diversity of bacteria and fungi in various tillage systems of sub-humid and semi-arid Kenya. Appl Soil Ecol 58:12–20.

32. Delmont TO, Prestat E, Keegan KP, Faubladier M, Robe P, Clark IM, Pelletier E, Hirsch PR, Meyer F, Gilbert JA, Le Paslier D, Simonet P, Vogel TM. 2012. Structure, fluctuation and magnitude of a natural grassland soil metagenome. ISME J 6:1677–1687.

33. Griffith GW, Bratton JH, Easton G. 2004. Charismatic megafungi - The conservation of waxcap grasslands. Br Wildl 16:31–43.

34. Griffith GW, Roderick K. 2008. Chapter 15 Saprotrophic basidiomycetes in grasslands: Distribution and function, p. 277–299. In British Mycological Society Symposia Series. Elsevier Ltd.

35. Silvertown J, Poulton P, Johnston E, Edwards G, Heard M, Biss PM. 2006. The Park Grass Experiment 1856-2006: Its contribution to ecology. J Ecol 94:801–814.

36. Seitzman BH, Ouimette A, Mixon RL, Hobbie EA, Hibbett DS. 2011. Conservation of biotrophy in hygrophoraceae inferred from combined stable isotope and phylogenetic analyses. Mycologia 103:280–290.

37. van der Heijden MGA, Martin FM, Selosse MA, Sanders IR. 2015. Mycorrhizal ecology and evolution: The past, the present, and the future. New Phytol 205:1406–1423.

38. Six J, Elliott ET, Paustian K. 2000. Soil macroaggregate turnover and microaggregate formation: A mechanism for C sequestration under no-tillage agriculture. Soil Biol Biochem 32:2099–2103.

39. Kleber M, Sollins P, Sutton R. 2007. A conceptual model of organo-mineral interactions in soils: Self-assembly of organic molecular fragments into zonal structures on mineral surfaces. Biogeochemistry 85:9–24.

40. Christensen BT. 2001. Physical fractionation of soil and structural and functional complexity in organic matter turnover. Eur J Soil Sci 52:345–353.

41. Chenu C, Stotzky G. 2002. Interactions between Microorganisms and Soil Particles: An Overview, p.. In Huang, PM, Bollag, J-M, Senesi, N (eds.), Interactions between Soil Particles and Microorganisms: Impact on the Terrestrial Ecosystem. John Wiley& Sons, Ltd.

42. Mills AL. 2003. Keeping in Touch: Microbial Life on Soil Particle Surfaces. Adv Agron 78:1–43.

43. Gleason FH, Crawford JW, Neuhauser S, Henderson LE, Lilje O. 2012. Resource seeking strategies of zoosporic true fungi in heterogeneous soil habitats at the microscale level. Soil Biol Biochem 45:79–88.

44. Sparrow FK. 1960. Aguatic phycomycetes. Second revised and enlarged edition.

45. Barr DJS. 2001. Chytridiomycota BT - Systematics and Evolution: Part A, p. 93–112. In McLaughlin, DJ, McLaughlin, EG, Lemke, PA (eds.),. Springer Berlin Heidelberg, Berlin, Heidelberg.

46. Gleason FH, Schmidt SK, Marano A V. 2010. Can zoosporic true fungi grow or survive in extreme or stressful environments ? Extremophiles 14:417–425.

47. Oades JM, Waters AG. 1991. Aggregate hierarchy in soils. Soil Res 29:815–828.

48. Golchin A, Oades JM, Skjemstad JO, Clarke P. 1994. Soil structure and carbon cycling. Soil Res 32:1043–1068.

49. Jastrow JD, Miller RM, Boutton TW. 1996. Carbon Dynamics of Aggregate-Associated Organic Matter Estimated by Carbon-13 Natural Abundance. Soil Sci Soc Am J 60:801–807.

50. Grundmann GL. 2004. Spatial scales of soil bacterial diversity - The size of a clone. FEMS Microbiol Ecol 48:119–127.

51. Miltner A, Bombach P, Schmidt-Brücken B, Kästner M. 2012. SOM genesis: microbial biomass as a significant source. Biogeochemistry 111:41–55.

52. Gregorich EG, Beare MH, Stoklas U, St-Georges P. 2003. Biodegradability of soluble organic matter in maize-cropped soils. Geoderma 113:237–252.

53. Balesdent J. 1996. The significance of organic separates to carbon dynamics and its modelling in some cultivated soils. Eur J Soil Sci 47:485–493.

54. John B, Yamashita T, Ludwig B, Flessa H. 2005. Storage of organic carbon in aggregate and density fractions of silty soils under different types of land use. Geoderma 128:63–79.

55. Constancias F, Prévost-Bouré NC, Terrat S, Aussems S, Nowak V, Guillemin JP, Bonnotte A, Biju-Duval L, Navel A, Martins JMF, Maron PA, Ranjard L. 2014. Microscale evidence for a high decrease of soil bacterial density and diversity by cropping. Agron Sustain Dev 34:831–840.

56. Freeman KR, Martin AP, Karki D, Lynch RC, Mitter MS, Meyera AF, Longcore JE, Simmons DR, Schmidt SK. 2009. Evidence that chytrids dominate fungal communities in high-elevation soils. Proc Natl Acad Sci U S A 106:18315–18320.

57. Benny GL, Humber RA, Voigt K. 2014. Zygomycetous Fungi: Phylum Entomophthoromycota and Subphyla Kickxellomycotina, Mortierellomycotina, Mucoromycotina, and Zoopagomycotina BT - Systematics and Evolution: Part A, p. 209–250. In McLaughlin, DJ, Spatafora, JW (eds.),. Springer Berlin Heidelberg, Berlin, Heidelberg.

58. Hemkemeyer M, Christensen BT, Tebbe CC, Hartmann M. 2019. Taxon-specific fungal preference for distinct soil particle size fractions. Eur J Soil Biol 94:103103.

59. Bonfante P, Venice F. 2020. Mucoromycota: going to the roots of plant-interacting fungi. Fungal Biol Rev 34:100–113.

60. Flemming HC, Wingender J. 2010. The biofilm matrix. Nat Rev Microbiol 8:623–633.

61. SevinçŞengör S. 2019. Review of Current Applications of Microbial Biopolymers in Soil and Future Perspectives, p. 275–299. In ACS Symposium Series. chapter, American Chemical Society.

62. Boutard M, Cerisy T, Nogue PY, Alberti A, Weissenbach J, Salanoubat M, Tolonen AC. 2014. Functional Diversity of Carbohydrate-Active Enzymes Enabling a Bacterium to Ferment Plant Biomass. PLoS Genet 10:e1004773.

63. Ranjard L, Nazaret S, Gourbière F, Thioulouse J, Linet P, Richaume A. 2000. A soil microscale study to reveal the heterogeneity of Hg(II) impact on indigenous bacteria by quantification of adapted phenotypes and analysis of community DNA fingerprints. FEMS Microbiol Ecol 31:107–115.

64. Mummey D, Holben W, Six J, Stahl P. 2006. Spatial stratification of soil bacterial populations in aggregates of diverse soils. Microb Ecol 51:404–411.

65. Davinic M, Fultz LM, Acosta-Martinez V, Calderón FJ, Cox SB, Dowd SE, Allen VG, Zak JC, Moore-Kucera J. 2012. Pyrosequencing and mid-infrared spectroscopy reveal distinct aggregate stratification of soil bacterial communities and organic matter composition. Soil Biol Biochem 46:63–72.

66. Neumann D, Heuer A, Hemkemeyer M, Martens R, Tebbe CC. 2013. Response of microbial communities to long-term fertilization depends on their microhabitat. FEMS Microbiol Ecol 86:71–84.

67. Hemkemeyer M, Pronk GJ, Heister K, Kögel-Knabner I, Martens R, Tebbe CC. 2014. Artificial soil studies reveal domain-specific preferences of microorganisms for the colonisation of different soil minerals and particle size fractions. FEMS Microbiol Ecol 90:770–782.

68. Poll C, Thiede A, Wermbter N, Sessitsch A, Kandeler E. 2003. Micro-scale distribution of microorganisms and microbial enzyme activities in a soil with long-term organic amendment. Eur J Soil Sci 54:715–724.

69. Mummey DL, Stahl PD. 2004. Analysis of Soil Whole- and Inner-Microaggregate Bacterial Communities. Microb Ecol 48:41–50.

70. Hemkemeyer M, Dohrmann AB, Christensen BT, Tebbe CC. 2018. Bacterial preferences for specific soil particle size fractions revealed by community analyses. Front Microbiol 9:1–13.

71. Sahariah P, Másson M. 2017. Antimicrobial Chitosan and Chitosan Derivatives: A Review of the Structure-Activity Relationship. Biomacromolecules 18:3846–3868.

72. Elsoud MMA, Kady EM El. 2019. Current trends in fungal biosynthesis of chitin and chitosan. Bull Natl Res Cent 43:1–12.

73. De Oliveira Franco L, Maia RDCC, Porto ALF, Messias AS, Fukushima K, De Campos-Takaki GM. 2004. Heavy metal biosorption by chitin and chitosan isolated from Cunninghamella elegans (IFM 46109). Brazilian J Microbiol 35:243–247.

74. Kaczmarek MB, Struszczyk-Swita K, Li X, Szczęsna-Antczak M, Daroch M. 2019. Enzymatic modifications of chitin, chitosan, and chitooligosaccharides. Front Bioeng Biotechnol 7:243.

75. Wan Ngah WS, Teong LC, Hanafiah MAKM. 2011. Adsorption of dyes and heavy metal ions by chitosan composites: A review. Carbohydr Polym 83:1446–1456.

76. Mueller B. 2015. Experimental Interactions Between Clay Minerals and Bacteria: A Review. Pedosphere 25:799–810.

77. Pankratov TA, Tindall BJ, Liesack W, Dedysh SN. 2007. Mucilaginibacter paludis gen. nov., sp. nov. and Mucilaginibacter gracilis sp. nov., pectin-, xylan and laminarin-degrading members of the family Sphingobacteriaceae from acidic Sphagnum peat bog. Int J Syst Evol Microbiol 57:2349–2354.

78. Urai M, Aizawa T, Nakagawa Y, Nakajima M, Sunairi M. 2008. Mucilaginibacter kameinonensis sp., nov., isolated from garden soil. Int J Syst Evol Microbiol 58:2046–2050.

79. Fredendall RJ, Stone JL, Pehl MJ, Orwin PM. 2020. Transcriptome profiling of Variovorax paradoxus EPS under different growth conditions reveals regulatory and structural novelty in biofilm formation. Access Microbiol 2:1–11.

80. Fish KE, Boxall JB. 2018. Biofilm microbiome (re)growth dynamics in drinking water distribution systems are impacted by chlorine concentration. Front Microbiol 9:1–21.

81. Andrews JS, Rolfe SA, Huang WE, Scholes JD, Banwart SA. 2010. Biofilm formation in environmental bacteria is influenced by different macromolecules depending on genus and species. Environ Microbiol 12:2496–2507.

82. Pantanella F, Berlutti F, Passariello C, Sarli S, Morea C, Schippa S. 2007. Violacein and biofilm production in Janthinobacterium lividum. J Appl Microbiol 102:992–999.

83. Yamada Y, Fukuda H, Inoue K, Kogure K, Nagata T. 2013. Effects of attached bacteria on organic aggregate settling velocity in seawater. Aquat Microb Ecol 70:261–272.

84. Li X, Yuan Y. 2002. Settling velocities and permeabilities of microbial aggregates. Water Res 36:3110–3120.

85. Pfeifer F. 2012. Distribution, formation and regulation of gas vesicles. Nat Rev Microbiol 10:705–715.

86. Walsby AE. 1972. Structure and function of gas vacuoles. Bacteriol Rev 36:1–32.

87. Guennoc CM, Rose C, Labbé J, Deveau A. 2018. Bacterial biofilm formation on the hyphae of ectomycorrhizal fungi: A widespread ability under controls? FEMS Microbiol Ecol 94:1–14.

88. Yurkov AM. 2018. Yeasts of the soil - obscure but precious. Yeast 35:369–378.

89. Henry S, Bru D, Stres B, Hallet S, Philippot L. 2006. Quantitative detection of the nosZ gene, encoding nitrous oxide reductase, and comparison of the abundances of 16S rRNA, narG, nirK, and nosZ genes in soils. Appl Environ Microbiol 72:5181–5189.

90. Bru D, Ramette A, Saby NPA, Dequiedt S, Ranjard L, Jolivet C, Arrouays D, Philippot L. 2011. Determinants of the distribution of nitrogen-cycling microbial communities at the landscape scale. ISME J 5:532–542.

91. Čuhel J, Šimek M, Laughlin RJ, Bru D, Chèneby D, Watson CJ, Philippot L. 2010. Insights into the effect of soil pH on N2O and N2 emissions and denitrifier community size and activity. Appl Environ Microbiol 76:1870–1878.

92. Di HJ, Cameron KC, Podolyan A, Robinson A. 2014. Effect of soil moisture status and a nitrification inhibitor, dicyandiamide, on ammonia oxidizer and denitrifier growth and nitrous oxide emissions in a grassland soil. Soil Biol Biochem 73:59–68.

93. Philippot L, Bru D, Saby NPA, Čuhel J, Arrouays D, Šimek M, Hallin S. 2009. Spatial patterns of bacterial taxa in nature reflect ecological traits of deep branches of the 16S rRNA bacterial tree. Environ Microbiol 11:3096–3104.

94. Miller MN, Zebarth BJ, Dandie CE, Burton DL, Goyer C, Trevors JT. 2009. Denitrifier community dynamics in soil aggregates under permanent grassland and arable cropping systems. Soil Sci Soc Am J 73:1843–1851.

95. Stienstra AW, Klein Gunnewiek P, Laanbroek HJ. 1994. Repression of nitrification in soils under a climax grassland vegetation. FEMS Microbiol Ecol 14:45–52.

96. Leininger S, Urich T, Schloter M, Schwark L, Qi J, Nicol GW, Prosser JI, Schuster SC, Schleper C. 2006. Archaea predominate among ammonia-oxidizing prokaryotes in soils. Nature 442:806–809.

97. Adair KL, Schwartz E. 2008. Evidence that ammonia-oxidizing archaea are more abundant than ammonia-oxidizing bacteria in semiarid soils of northern Arizona, USA. Microb Ecol 56:420–426.

98. de Vries W, Leip A, Reinds GJ, Kros J, Lesschen JP, Bouwman AF. 2011. Comparison of land nitrogen budgets for European agriculture by various modeling approaches. Environ Pollut 159:3254–3268.

99. Fowler D, Coyle M, Skiba U, Sutton MA, Cape JN, Reis S, Sheppard LJ, Jenkins A, Grizzetti B, Galloway JN, Vitousek P, Leach A, Bouwman AF, Butterbach-Bahl K, Dentener F, Stevenson D, Amann M, Voss M. 2013. The global nitrogen cycle in the Twentyfirst century. Philos Trans R Soc B Biol Sci 368:20130164.

100. Cuadros J. 2017. Clay minerals interaction with microorganisms: a review. Clay Miner 52:235–261.

101. Norton J, Ouyang Y. 2019. Controls and adaptive management of nitrification in agricultural soils. Front Microbiol 10:1–18.

102. Deveau A, Bonito G, Uehling J, Paoletti M, Becker M, Bindschedler S, Hacquard S, Hervé V, Labbé J, Lastovetsky OA, Mieszkin S, Millet LJ, Vajna B, Junier P, Bonfante P, Krom BP, Olsson S, van Elsas JD, Wick LY. 2018. Bacterial-fungal interactions: Ecology, mechanisms and challenges. FEMS Microbiol Rev 42:335–352.

103. Sharma K, Palatinszky M, Nikolov G, Berry D, Shank EA. 2020. Transparent soil microcosms for live-cell imaging and non-destructive stable isotope probing of soil microorganisms. Elife 9:1–28.

104. Kohlmeier S, Smits THM, Ford RM, Keel C, Harms H, Wick LY. 2005. Taking the fungal highway: Mobilization of pollutant-degrading bacteria by fungi. Environ Sci Technol 39:4640–4646.

105. Fomina M, Skorochod I. 2020. Microbial interaction with clay minerals and its environmental and biotechnological implications. Minerals 10:1–54.

106. Navel A, Martins JMF. 2013. Effect of long term organic amendments and vegetation of vineyard soils on the microscale distribution and biogeochemistry of copper. Sci Total Environ 466-467:681–689.

107. Hood-Nowotny R, Hinko-Najera Umana N, Inselbacher E, Oswald-Lachouani P, Wanek W. 2010. Alternative methods for measuring inorganic, organic, and total dissolved nitrogen in soil. Soil Sci Soc Am J 74:1018–1027.

108. Ranjard L, Brothier E, Nazaret S. 2000. Sequencing bands of ribosomal intergenic spacer analysis fingerprints for characterization and microscale distribution of soil bacterium populations responding to mercury spiking. Appl Environ Microbiol 66:5334–5339.

109. Bokulich NA, Mills DA. 2013. Improved selection of internal transcribed spacerspecific primers enables quantitative, ultra-high-throughput profiling of fungal communities. Appl Environ Microbiol 79:2519–2526.

110. Core R Team. 2019. A Language and Environment for Statistical Computing. R Found Stat Comput. R Foundation for Statistical Computing.

111. Schmieder R, Edwards R. 2011. Quality control and preprocessing of metagenomic datasets. Bioinformatics 27:863–864.

112. Cole JR, Wang Q, Fish JA, Chai B, McGarrell DM, Sun Y, Brown CT, Porras-Alfaro A, Kuske CR, Tiedje JM. 2014. Ribosomal Database Project: Data and tools for high throughput rRNA analysis. Nucleic Acids Res 42:633–642.

113. Katoh K, Misawa K, Kuma KI, Miyata T. 2002. MAFFT: A novel method for rapid multiple sequence alignment based on fast Fourier transform. Nucleic Acids Res 30:3059–3066.

114. Price MN, Dehal PS, Arkin AP. 2009. Fasttree: Computing large minimum evolution trees with profiles instead of a distance matrix. Mol Biol Evol 26:1641–1650.

